# DeepParcellation: a novel deep learning method for robust brain magnetic resonance imaging parcellation in older East Asians

**DOI:** 10.1101/2022.04.06.487283

**Authors:** Eun-Cheon Lim, Uk-Su Choi, Kyu Yeong Choi, Jang Jae Lee, Yul-Wan Sung, Seiji Ogawa, Byeong Chae Kim, Kun Ho Lee, Jungsoo Gim, the Alzheimer’s Disease Neuroimaging Initiative

**Affiliations:** Gwangju Alzheimer’s & Related Dementia Cohort Research Center, Chosun University, Gwangju 61452, Republic of Korea; BK FOUR Department of Integrative Biological Sciences, Chosun University, Gwangju 61452, Republic of Korea; Department of Biomedical Science, Chosun University, Gwangju 61452, Republic of Korea; Korea Brain Research Institute, Daegu 41062, Republic of Korea; Neurozen Inc., Seoul 06236, Republic of Korea; Kansei Fukushi Research Institute, Tohoku Fukushi University, Sendai, Miyagi 9893201, Japan; Department of Neurology, Chonnam National University Medical School, Gwangju 61469, Republic of Korea

**Keywords:** Deep learning, brain, 3D magnetic resonance imaging, DeepParcellation, parcellation

## Abstract

Accurate parcellation of cortical regions is crucial for distinguishing morphometric changes in aged brains, particularly in degenerative brain diseases. Normal aging and neurodegeneration precipitate brain structural changes, leading to distinct tissue contrast and shape in people aged > 60 years. Manual parcellation by trained radiologists can yield a highly accurate outline of the brain; however, analyzing large datasets is laborious and expensive. Alternatively, newly-developed computational models can quickly and accurately conduct brain parcellation, although thus far only for the brains of Caucasian individuals. DeepParcellation, our novel deep learning model for 3D magnetic resonance imaging (MRI) parcellation, was trained on 5,035 brains of older East Asians (Gwangju Alzheimer’s & Related Dementia) and 2,535 brains of Caucasians. We trained full 3D models for N-way individual regions of interest using memory reduction techniques. Our method showed the highest similarity and robust reliability among age-ethnicity groups, especially when parcellating the brains of older East Asians.

## 1. Introduction

Population growth, in association with aging, is a driving force for the increasing incidence of neurodegenerative diseases. Brain aging is reflected in structural changes and functional decline of the brain [*1*]. Estimating the brain’s biological age and monitoring the progression of age-related diseases [*2*] demand accurate brain parcellation methods. However, earlier parcellation methods have overlooked aging morphology and ethnic differences, raising several concerns.

The first concern is that the brains of people aged > 60 years show robustly different region-specific patterns compared with those of younger individuals (20s– 40s). For instance, the contrast between gray matter (GM) and white matter (WM) is usually higher in younger brains than in older brains because of changes in the amount of water in GM and WM tissues driven by myelin structural changes [*3*]. Subcortical structures show heterogeneous T1 and T2* values across regions due to changes in the composition of myelin and iron [*4*]. Abnormal volume and shape changes in the brain of older persons are observed as ventricular enlargement [*5*], WM hyperintensities [*6*], WM/GM atrophy [*7*], and heterogeneous subcortical brain volumes [*4*].

The second concern is that brain volume and shape differ between East Asians and Caucasians [*8*]. Brains of Japanese individuals, for example, show morphological differences in the inferior parietal lobes, occipital regions, and posterior temporal regions compared with those of Europeans. The overall notion is that the brains of Japanese participants are shorter and wider than those of European participants [*9*]. A recent study validated the interethnic differences in cortical volume, cortical thickness, cortical surface area, and GM intensities [*10*], and reported that the brains of Chinese participants showed larger structural aspects in the temporal lobe and cingulate gyrus, but smaller ones in the parietal and frontal lobes than the brains of Caucasian individuals.

The final concern is related to computation time. There is a growing interest in collecting and studying brain magnetic resonance imaging (MRI) cohort data of East Asian individuals [*11*]. A fast and reliable segmentation method is critical for such studies because conventional methods require long computation times to improve accuracy [*12*]. This can take many hours per brain, depending on computing power or algorithm complexity. Although recent advances have reduced the computation time, they are still not sufficient to handle big cohort data. To overcome these performance issues, deep learning approaches have recently been considered as a suitable solution in the neuroimaging field [*13*–*18*]. However, these models may not be directly applicable to brain parcellation of older East Asian individuals in terms of their runtime and accuracy, as they are based on brains of Caucasian individuals.

Deep learning models for brain segmentation and parcellation usually suffer from a tradeoff between the image dimensions and memory requirements. Based on image dimensions, models can be divided into four categories: 2D, 2.5D, partial 3D, and full 3D models. 2D models are the simplest, whereby only a single slice is segmented [*13*]. They lose 3D contexts orthogonal to the selected plane and do not provide an aggregated 3D view of the parcellated regions of interest (ROIs). In contrast, 2.5D models attempt to reconstruct a 3D view from slice-wise segmentations [*19*]. This strategy could reduce some inconsistencies between slices by considering adjacent contexts. However, the aggregation could still create artifacts at random positions, degrading the overall accuracy [*19*]. Partial 3D models are the most common and use partial 3D images/patches derived from the whole image either by sub-or down-sampling. Partial 3D-based models can usually observe local 3D contexts, producing better parcellation for a certain area, while losing some global contexts [*14, 17, 18*]. A notable exception to this limitation is a cascaded model that can capture both global and local contexts by using down-sampled and cropped images of the original resolution. However, this strategy is not capable of handling ROIs of varying sizes. Full 3D models can capture 3D contexts, intrinsically reducing inconsistency between slices and, in turn, potentially yielding high accuracy [*15, 16*]. However, the memory requirements become intractable, owing to the required increases in model parameters.

ROI number and size usually govern a model’s performance, mainly due to class imbalance. A model may predict a few ROIs of larger volumes with higher accuracy and shorter computing time than those segmenting several smaller ROIs. A few early models focused only on a single ROI, such as the hippocampus [*17*]. Next-tier models can parcellate three representative tissues, including GM, WM, and cerebrospinal fluid [*20*], while finer-grained models collocate ROI predictions in the left and right cerebral hemispheres. Of particular interest is SkipDeconv-Net (SD-Net) [*15*], which adopted UNet [*21*] and DeconvNet [*15*]. The SD-Net author introduced error corrective boosting (ECB), which updates high weights for classes with low accuracy per epoch, giving increased attention to those classes. However, ECB was applied only to the weighted cross-entropy and not to dice loss. A pioneering model handling > 50 ROIs, which can segment MR slices into 56 classes using a 2D convolutional neural network (CNN), was introduced in 2011 [*13*]. Some groups claimed to have successfully created a model that can parcellate > 90 ROIs [*18, 19*]. NeuroNet was trained on large-scale samples (N = 5,723) from the UK Biobank imaging study, and used three different segmentation tools, FSL, SPM, and MALP-EM, to generate label maps from T1-weighted images [*16*]. However, there are some limitations to NeuroNet, such as the image dimensions of 128 × 128 × 128 and the difficulty to improve the low accuracy of some ROIs unless training is performed with weighted losses. Recently, FastSurferCNN achieved state-of-the-art performance in the parcellation of 95 ROIs using the 2.5D UNet with competitive dense blocks [*19*]. Notably, only a few models were trained and tested on > 500 subjects in which the number of subjects is a very important factor in testing the reliability and validity of a given model [*16, 18, 19*]. A summary of the other available models is provided in Tables A.1-A.4.

Accurate brain segmentation and parcellation are necessary for acquiring precise quantitative values of brain regions, including volume and cortical thickness[*8, 22, 23*]. These measurements have been used for brain age prediction [*1*] and as biomarkers for neurodegenerative diseases including Parkinson’s disease (PD) [*24*] and Alzheimer’s disease (AD) [*25*].

In this study, we propose a novel 3D deep learning model, DeepParcellation, focusing on the brains of older East Asian individuals, which can parcellate 109 ROIs based on the Desikan-Killiany-Tourville (DKT) atlas. Our model employs 3D UNet architectures combined with inception blocks, dilated convolutions, and attention gates. The proposed model was robustly evaluated in (1) similarity of parcellated regions using dice coefficient (DICE), averaged Hausdorff Distance (aHD), (2) intra-subject reliability using the intra-class correlation coefficient (ICC), and (3) between-group variability between cognitively normal (CN) people and patients with AD.

## 2. Materials and methods

All participants provided informed consent in accordance with the institutional review board of Chosun University Hospital, Republic of Korea.

### 2.1. Experimental Design

The primary aim of the study was to provide robust brain features for downstream analyses in studies of neurodegenerative diseases, aging, and biomarkers for monitoring patients in follow-ups. To enable support for unlimited number of ROIs, we introduced the N-way-weight strategy. Following the divide-and-conquer concept, we performed individual training for each ROI, avoiding competition during training so that ROI weights are independent. In addition, we integrated three memory reduction techniques to overcome a limitation in computational resources while retaining the full 3D characteristics: inception blocks, dilated convolutions, and attention gates.

To evaluate model performance, we collected brain MRI data of people of East Asian and Caucasian origins. We focused on similarity and robustness measures by which the accuracy and robustness of the downstream analyses could be improved.

### 2.2. Model background

UNet was initially developed for segmenting 2D biomedical images [*21*]. UNet follows an encoder–decoder structure for unsupervised learning, where the encoder (contracting path) captures global contexts, while the decoder (expanding path) performs detailed localizations. Skip connections in the expanding path combine contextual information and spatial locations.

A deep learning model can improve accuracy through a deeper or wider network structure. However, these structural changes lead to an increase in model capacity, causing overfitting and gradient vanishing problems. The Inception (or GoogLeNet) block mitigates these problems by introducing 1 × 1 or 1 × 1 × 1 convolution blocks, which reduce the number of feature maps while increasing depth [*32*].

In a convolutional layer, the kernel size determines the receptive field area, which represents the feature size. Multi-scale kernels can improve model performance, but also increase the number of parameters. Dilated convolution was introduced to enable larger receptive fields while maintaining the same number of parameters [*33*]. For instance, for a CNN with a kernel of size 3 × 3 and a dilation rate of 2, the receptive field becomes 5 × 5 while keeping the number of parameters to nine because every second row and column of the field will be skipped.

Attention is a mechanism that focuses more on features relevant to the target than on those less relevant [*34*]. Soft attention keeps attention on the global context, while hard attention observes a partial context, such as patches of an input. Soft attention can be implemented as a skip connection in UNet [*35*].

We developed DeepParcellation using N-way multiple 3D UNet architectures combined with inception blocks, dilated convolutions, and attention gates (Fig. 1)

**Fig. 1.**
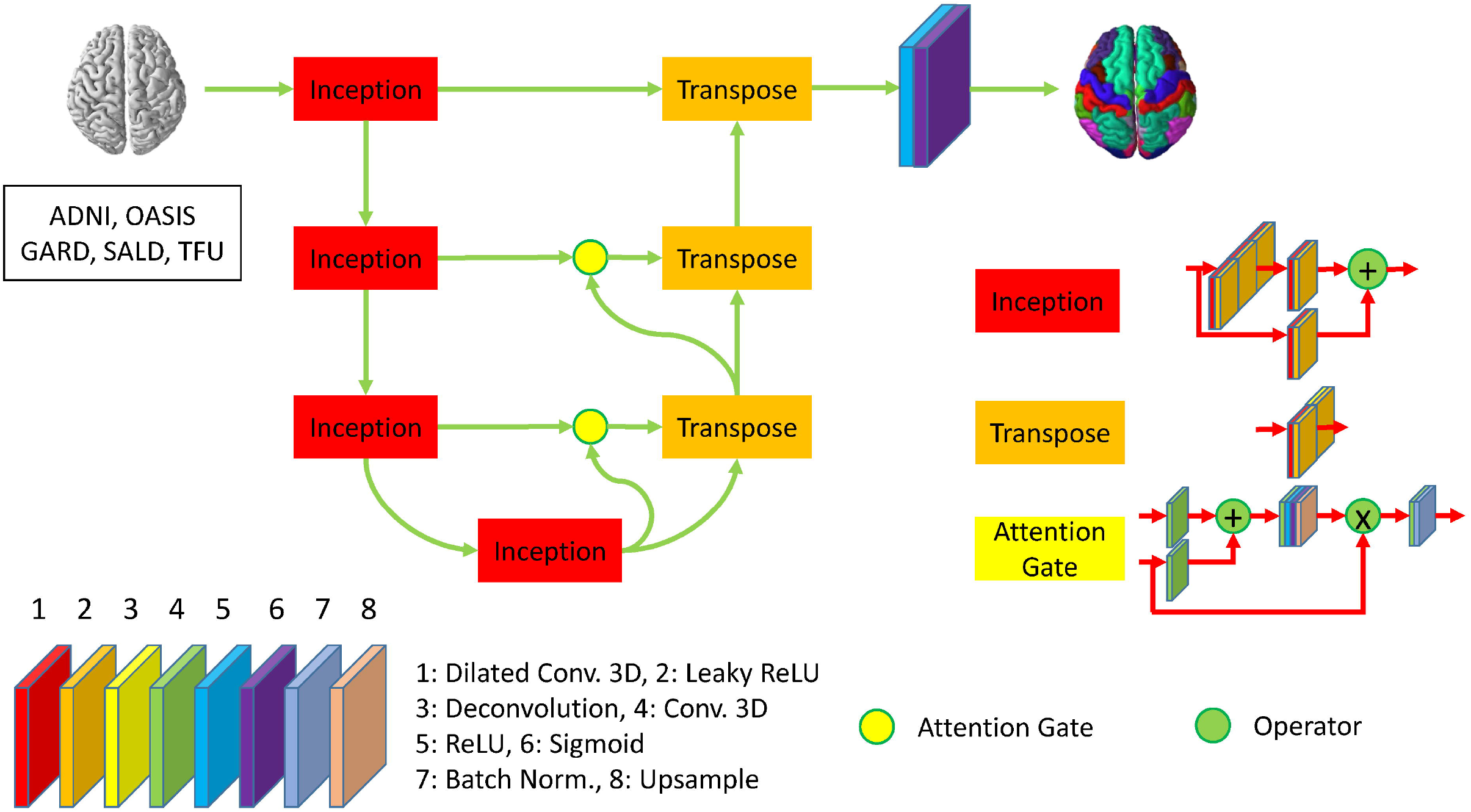
Description of DeepParcellation network architecture for a single region of interest. The network includes four inception blocks and three transpose blocks which consist of eight different layers. The second, third, and fourth inception blocks are connected to transpose blocks through attention gates. The last transpose block is activated through ReLU and Sigmoid function to predict parcellated brain image.

### 2.3. Datasets

#### 2.3.1. East Asian old age (OA) dataset

MR images of 5,035 older Koreans, aged > 50 years, were collected from the Gwangju Alzheimer’s & Related Dementia (GARD) dataset (Table 1). GARD data were divided into 4,028, 503, and 504 subjects for training, validation, and test sets, respectively (Table A.6). MRI data were acquired using 3.0 T (Skyra, Siemens, Munich, Germany) scanners. T1-MPRAGE sequences were acquired with the following parameters: repetition time (TR) = 2,300 ms, echo time (TE) = 2.14 ms, inversion time (TI) = 900 ms, field of view (FOV) = 256 × 256, and voxel size = 0.8 × 0.8 × 0.8 mm^3^. T2-SPACE sequence was acquired with the following parameters: TR = 2,300 ms, TE = 2.143 ms, TI = 900 ms, FOV = 256 × 256, and voxel size = 0.8 × 0.8 × 0.8 mm^3^.

**Table 1.** Characteristics of study samples. The distribution of age, sex, and sample size are described.

| Dataset | No. of Subjects | Age | Male (%) | Female (%) |
| --- | --- | --- | --- | --- |
| GARD | 5035 (Sex unspecified: 51) | 35 - 100 (72.62±0.15) | 40.62 | 58.37 |
| SALD | 487 (Sex unspecified: 2) | 19 - 80 (45.31±1.55) | 36.76 | 62.83 |
| TFU | 140 | 18 - 22 (19.05±0.14) | 60.00 | 40.00 |
| ADNI | 956 | 55 - 97 (74.25±0.46) | 50.21 | 49.79 |
| OASIS | 1579 | 18 - 98 (68.34±0.68) | 42.62 | 57.38 |

Abbreviations: ADNI, Alzheimer's Disease Neuroimaging Initiative; GARD, Gwangju Alzheimer's & Related Dementia; OASIS, Open Access Series of Imaging Studies; SALD, Southwest University Adult Lifespan Dataset; TFU, Tohoku Fukushi University

We used MR images of 116 Chinese individuals aged > 60 years from the Southwest University Adult Lifespan Dataset (SALD) only for model evaluation. Details of the MRI protocol are described in the study by Wei et al. [*27*].

#### 2.3.2. East Asian young age (YA) dataset

MR images of 140 young Japanese individuals (mean age: 19.05, 18–22 years) were collected from the Tohoku Fukushi University (TFU) dataset [*36*]. TFU MRI data were acquired using 3.0 T (Skyra, Siemens) scanners. T1-MPRAGE sequences were acquired with the following parameters: TR = 1,900 ms, TE = 2.52 ms, TI = 900 ms, FOV = 256 × 256, and voxel size = 1 × 1 × 1 mm^3^.

MR images of 154 young Chinese individuals < 30 years from the SALD were used for model evaluation.

#### 2.3.3. Caucasian OA dataset

MR images of 75 older Caucasians, aged > 60 years, were collected from the Alzheimer’s Disease Neuroimaging Initiative (ADNI) dataset. Details of the dataset are described on the ADNI website (http://adni.loni.usc.edu).

MR images of 149 older Caucasians, aged > 60 years, were collected from the Open Access Series of Imaging Studies (OASIS) dataset. Details of the dataset are described on the OASIS website (https://www.oasis-brains.org).

#### 2.3.4. Caucasian YA dataset

MR images of 107 young Caucasians < 30 years were collected from the OASIS dataset. Details of the dataset are described on the OASIS website (https://www.oasis-brains.org).

#### 2.3.5. Intra-subject reliability dataset

To evaluate intra-subject reliability among ethnicities, we used three young Japanese subjects with six repeated acquisitions [*36*] and three Caucasian subjects with 40-times repeated acquisition within 31 days [*37*].

### 2.4. Preparation for model training

The labeled images were reconstructed using T1 and T2 images to reduce topological mismatches when T2 images were available. To run the *recon-all* command with the T2 argument, T2-SPACE images were spatially registered to the T1 space. We adopted the rigid-body registration strategy to minimize registration errors, because T1-MPRAGE and T2-SPACE were acquired from the same scanner. The registered T2-SPACE and T1-MPRAGE images were used as inputs for running the FreeSurfer *recon-all* procedure with the following commands:

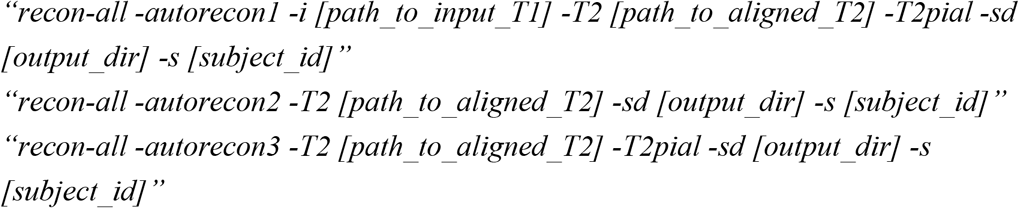

### 2.5. Model training

We did not perform any data augmentation, which is very common for studies with a limited number of samples. We initially pre-trained 112 independent ROIs defined in automatic cortical parcellation and automatic segmentation volume with different numbers of rounds consisting of multiple epochs using Keras [*38*] packages, mainly using six Tesla V-100 GPUs (graphics processing units) with 16 GB memory, and partially using six Tesla A-40 with 48 GB memory. Then, we performed transfer learning on 101 ROIs defined in the DKT atlas. Transfer learning is a learning strategy that reuses parts (or the whole) of the knowledge gained in previous tasks on a different but related task. Each epoch took approximately 1 h for 5,392 MRIs of a single ROI, and the model was trained for 121 days. The loss function improved curves near segmentation boundaries by using the DICE [*39*], voxel classification accuracy by assigning more weights to ROI masks according to class frequency between background and ROI voxels, and by using binary cross-entropy. We minimized the combined loss of binary cross-entropy and DICE using the Adam optimizer [*40*]. The learning rate was fixed to 0.0001, and the seed for the random number generator and optimizer weights were reset per round to overcome the local minima problem. Details of the training epoch information are shown in Fig. A.1.

### 2.6. Aggregation of individual predictions

All N-ROI masks predicted by DeepParcellation require an aggregation step, which provides an integrated view of individual probability maps. A single predicted result represents a probability map of the ROI voxels after passing the input to the sigmoid function. Because all probability maps share identical dimensions, we can determine the most likely classes of every voxel in 3D coordinates. Given input ROIs, x, we calculate the final probability map using the softmax function (σ) as follows:

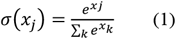

where, input probability vector *x* = {*x*_0_, *x*_1_, …, *x*_*n*−1_}, and *k* is the number of ROIs.

### 2.7. Statistical Analysis

#### 2.7.1. Similarity

DICE is a metric for evaluating segmentation accuracy. Given a binary mask of ground truth T and prediction P (voxels of the given class marked with 1 and background with 0), the DICE is defined as follows:

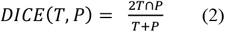

The highest value for DICE is 1, which represents a situation when T and P are perfectly matched. DICE is a widely accepted metric because it allows direct observation of the similarity between T and P. However, DICE may not capture the variability in fundi of different sulci (or simply the curvature) around ROIs’ boundaries.

Yet another metric, the Hausdorff distance (HD), can be used to measure how far two surfaces are from each other, bridging the gap in DICE. Given ground truth G and segmentation S, the HD is defined as follows:

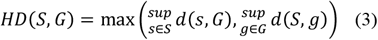

where sup represents the supremum or the greatest lower bound and is 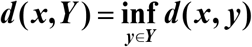 the distance ***x*** ∈ ***S*** from a point to the subset ***Y*** ⊆ ***S***.

Alternatively, the supremum distance or directed HD can be denoted as follows:

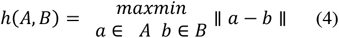

where, norm (||) is the Euclidean distance. However, directed HD is prone to being affected by noise and outliers; therefore, we could take the aHD. To calculate aHD, we replace distance as follows:

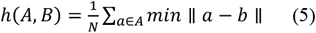

Equivalently, we can use a simplified equation of aHD as follows:

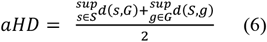

We calculated DICE, and aHD for the different age groups (OA and YA) of East Asians and Caucasians by comparing the predicted masks with the outputs of FreeSurfer (ground truth).

To clearly observe metric differences between our proposed model and another method, FreeSurfer, we calculated fold changes using the mean measurements of the other method as the baseline, and Cohen’s d. Given two groups, Cohen’s d is calculated as follows:

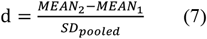

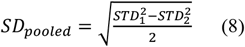

where, MEAN is the mean of a group and STD is a standard deviation of a group.

#### 2.7.2. Intra-subject reliability

For test-retest reliability evaluation, we adopted ICC (2, *k*) with a two-way random-effects model [*41*]. The definition of ICC (2, *k*) is that randomly selected *k* raters rate each target, and the reliability is estimated for the average of *k* ratings. Thus, we defined repeated measurements of the number of voxels as raters and subjects as targets. We calculated ICCs using the test-retest dataset for brain volume measurement, with 18 MRI scans from three young Japanese subjects and 120 MRI scans from three young Caucasian subjects [*37*]. ICCs were calculated using the voxel count of each ROI mask given by FreeSurfer and the proposed model.

#### 2.7.3. Between-group variability evaluation

To evaluate the sensitivity to inter-group variations, we compared the normalized cortical volume of each ROI mask given by FreeSurfer and our model between East Asian CN and AD groups using the GARD dataset. The normalized volume was calculated by dividing the voxel number of individual ROIs by the total voxel counts of all parcellated regions. Independent t-tests and f-tests were conducted between groups.

## 3. Results

### 3.1. Similarity evaluation

We calculated the DICEs by comparing the predicted ROIs of DeepParcellation and FastSurfer with the outputs of FreeSurfer version 7.1 as surrogates for the ground truth. FastSurfer was trained using FreeSurfer version 6.0, in a way that direct DICE comparisons between DeepParcellation and FastSurfer are infeasible. In this comparison, FreeSurfer version 7.1 outputs are likely to be unseen data from the FastSurfer model’s perspective; thus, the DICEs of FastSurfer serve as baselines for calculating fold changes. In the OA group, the mean DICEs for all 101 of DeepParcellation’s ROIs were higher than those of FastSurfer (Fig. 2A). Similarly, higher mean DICE values were observed with DeepParcellation, except for two ROIs (left and right superior temporals) in the YA group. Specifically, FastSurfer showed a higher fold change than DeepParcellation only in the right superior temporal ROI with a large Cohen’s d value (Fig. 2A). In the comparisons between ethnic groups, DeepParcellation showed higher similarities for all ROIs in the Asian group, while FastSurfer showed higher similarities for 74 out of 101 ROIs in the Caucasian group. With Cohen’s d criteria, DeepParcellation showed higher fold changes than FastSurfer except for three ROIs (left and right white matters and right superior temporal). We could not calculate the DICEs of six ROIs (CC_Posterior, CC_Anterior, CC_Mid_Posterior, CC_Mid_Anterior, CC_Central, and Optic-Chiasm) with FastSurfer because it did not produce ROI predictions with the default option.

**Fig. 2.**
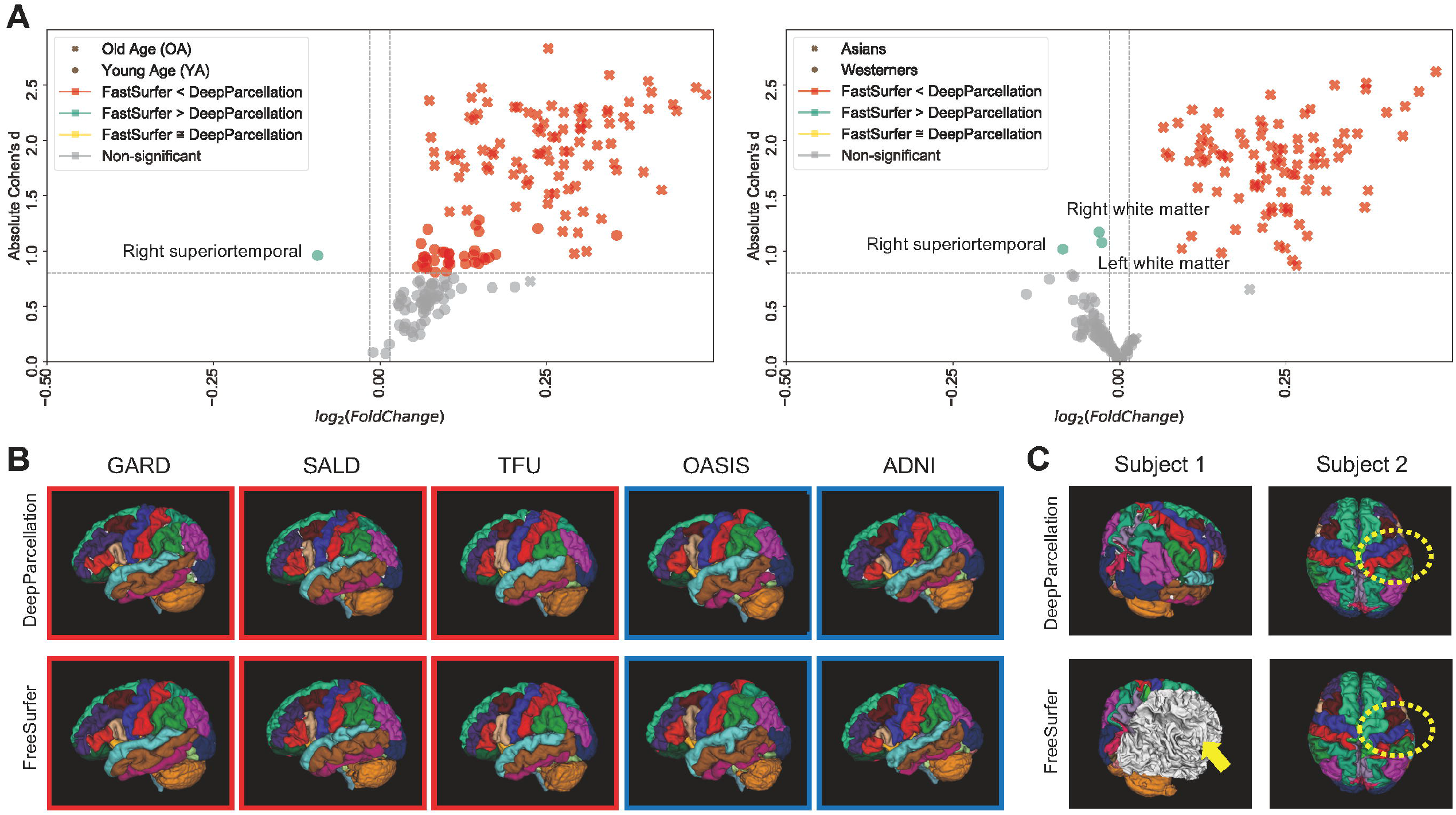
Performance of DeepParcellation. **(A)** Dice coefficient (DICE) comparison between DeepParcellation and FastSurfer. Using FreeSurfer output as surrogate for the ground truth, FastSurfer DICEs were baselines for calculating fold changes. Cohen’s d values were calculated using mean DICEs in two aspects (age and ethnicity). The horizontal line defines a Cohen’s d of 0.8, representing a large effect size. **(B)** Surface construction of parcellated brain images from representative subjects of different datasets. Red and blue squares indicate brains of Asians and Caucasians, respectively. (C) Examples of better parcellation with DeepParcellation compared with FreeSurfer. (a) Failures of right cortical parcellation in FreeSurfer (yellow arrows). (b) Wrong parcellation of right precentral and postcentral gyri in FreeSurfer (yellow dashed circle).

We also compared the DICEs among different groups using only DeepParcellation predictions to observe the effects of age and ethnicity. The performance of DeepParcellation was superior on brains of East Asians to that in brains of Caucasian individuals, and on OA compared to YA datasets (Fig. 3A). Among all groups, the overall highest average DICE (0.85) was observed in the East Asian OA group (one-way analysis of variance and post-hoc Tukey honestly significant difference, p < 0.001). In East Asians, the OA group showed a comparable number of brain regions with higher DICEs (43 out of 101 regions) to the YA group (Fig. 3A). The aHD of the East Asian OA group (0.28) was significantly lower than that of the East Asian YA group (0.30) (post-hoc, p < 0.001) (Fig. 3B) and Caucasian YA group (0.43) (post-hoc, p < 0.001). In addition, the aHD of the East Asian YA group (0.30) was significantly lower than that of the Caucasian YA group (0.43) (post-hoc, p < 0.001).

**Fig. 3.**
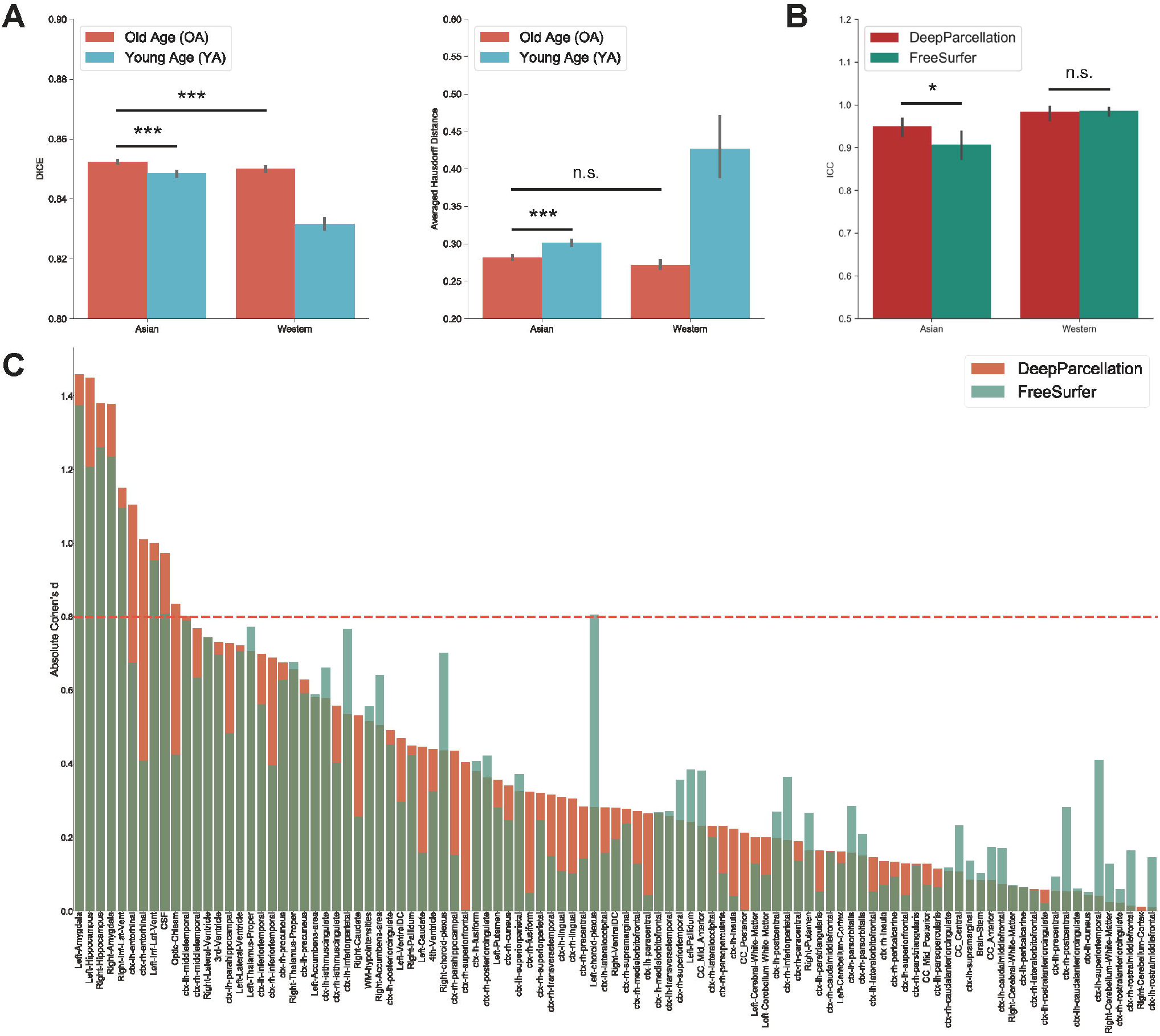
Evaluation of DeepParcellation. **(A)** Similarity evaluation among various age and ethnicity groups. (a) Dice coefficient (DICE) comparison. (b) aHD comparison between groups. Brains of older Asian individuals showed significantly higher DICE values compared with brains of young Asian individuals and older Caucasian individuals. Brains of older Asian individuals also showed significantly higher averaged Hausdorff Distance values compared with brains of young Asian individuals. **(B)** Intra-subject reliability evaluation between DeepParcellation and FreeSurfer across ethnicities. DeepParcellation showed a significantly higher intra-class correlation coefficient than FreeSurfer in brains of Asians. **(C)** Between-group variability evaluation using Cohen’s ds between DeepParcellation and FreeSurfer across parcellated regions of interest. The red dashed line indicates a Cohen’s d of 0.8 (large effect size). n.s. not significant; * significant at p < 0.05; ** significant at p < 0.01; *** significant at p < 0.001.

### 3.2. Intra-subject reliability

DeepParcellation showed a significantly higher average ICC (0.95) than that of FreeSurfer (0.91) in East Asians (Fig. 3B). There was no significant difference in DeepParcellation ICCs between East Asians and Caucasians (0.95 and 0.98, respectively), but significantly different FreeSurfer ICCs were observed between East Asians and Caucasians (0.91 and 0.99, respectively).

### 3.3. Between-group variability

In 61 out of 101 regions, DeepParcellation achieved higher statistical power of group differences between the CN and AD groups compared to FreeSurfer (Fig. 3C, and Fig. 4). We selected nine highly ranked regions sorted by the difference in the negative logarithm of p-values between CN and AD groups: bilateral entorhinal cortex, bilateral amygdala, bilateral hippocampus, and bilateral inferior lateral ventricles (Fig. 4).

**Fig. 4.**
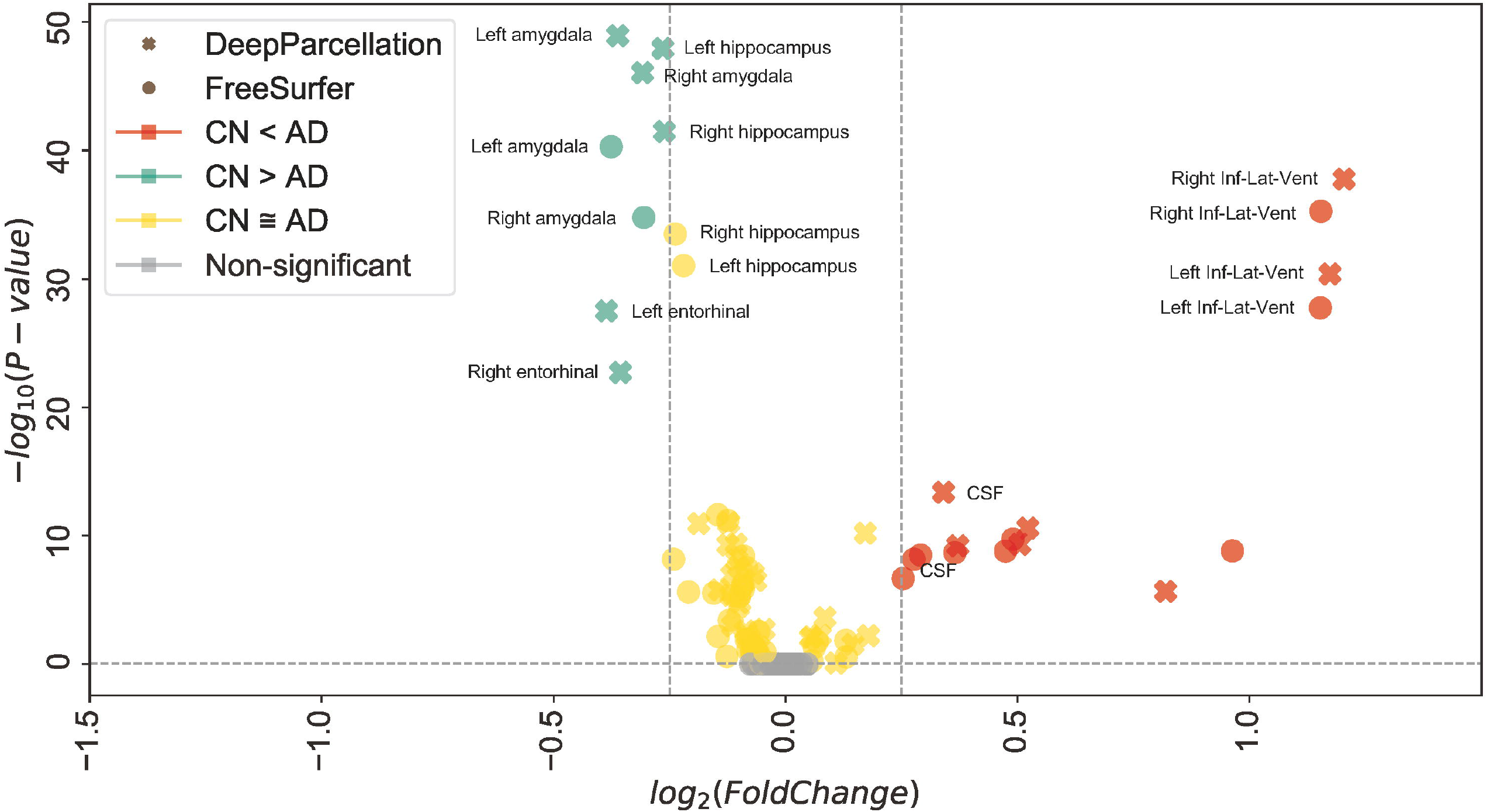
Robust dissociation of different diagnosed groups in DeepParcellation. Mean of normalized cortical volume with DeepParcellation (X) and FreeSurfer (O) in selected highly-ranked regions sorted by the difference in the negative logarithm of p-values between the cognitive normal (CN) and Alzheimer’s disease (AD) groups. DeepParcellation showed higher significance of normalized cortical volume differences than did FreeSurfer in regions highly associated with AD.

### 3.4. Processing success rate of DeepParcellation and FreeSurfer

We reported the success rate of DeepParcellation and FreeSurfer with default *recon-all* commands across different datasets (Table 2).

**Table 2.** Processing success rate of DeepParcellation and Freesurfer showing absolute numbers and percentage.

| **Dataset** | **Module** | **Count** | **Total** | **Success Rate (%)** |
| --- | --- | --- | --- | --- |
| ADNI | DeepParcellation | **4500** | 4502 | **99.96** |
| ADNI | FreeSurfer | 4461 | 4502 | 99.09 |
| OASIS | DeepParcellation | **5162** | 5162 | **100.00** |
| OASIS | FreeSurfer | 4889 | 5162 | 94.71 |
| GARD | DeepParcellation | **7337** | 7337 | **100.00** |
| GARD | FreeSurfer | **7337** | 7337 | **100.00** |
| SALD | DeepParcellation | **494** | 494 | **100.00** |
| SALD | FreeSurfer | 487 | 494 | 98.58 |
| TFU | DeepParcellation | **153** | 153 | **100.00** |
| TFU | FreeSurfer | 152 | 153 | 99.35 |

Abbreviations: ADNI, Alzheimer's Disease Neuroimaging Initiative; GARD, Gwangju Alzheimer's & Related Dementia; OASIS, Open Access Series of Imaging Studies; SALD, Southwest University Adult Lifespan Dataset; TFU, Tohoku Fukushi University

DeepParcellation failed only in two subjects in the ADNI dataset and succeeded in all other subjects in the other datasets. On the other hand, FreeSurfer failed in some subjects in all datasets (41, 273, 41, 7, and 1 subject in the ADNI, OASIS, GARD, SALD, and TFU datasets, respectively).

Most subjects were successfully processed by both DeepParcellation and FreeSurfer (Fig. 2B, and Fig. A.3A), but showed relatively lower average DICEs. We found that lower DICEs in FreeSurfer were due to failures in parcellations and misannotations of several brain regions (Fig. 2C, and Supplementary Fig. A.3B).

### 3.5. Runtime

We reported the runtime of DeepParcellation and FreeSurfer with the default *recon-all* command. DeepParcellation consistently performed full parcellation in about 2 min, at 30 s per sample, using a single GPU. The runtime of DeepParcellation using CPUs (central processing units) depends on the CPU number, but it did not improve when it was higher than the ROI number. The runtime of FreeSurfer fluctuated, with a median time of approximately 13 h and 9 h using a single CPU and 24 CPUs, respectively (Table A.5).

## 4. Discussion

Herein, we proposed a novel full 3D deep learning model for automatic brain MRI parcellation that shows comparable or better performance in terms of similarity and reliability for the brain of older East Asian individuals than an existing model, FreeSurfer. Previous deep learning models have utilized several tens or hundreds of samples of brains belonging to Caucasian individuals [*16, 18*], as East Asian cohort datasets were not sufficiently established or not publicly available, with the exception of a few cases [*26, 27*]. These East Asian studies consisted of hundreds of subjects, but the sample size may not be sufficient to train a deep learning model for older East Asian individuals. In contrast, the sample size (GARD, N = 7,166) and age range (mean age: 72.62, 35–100 years) of our study were suitable for implementing a deep learning model representing older East Asian individuals.

Our model was trained using the full 3D context. This forgoes the need for aggregating contexts of three orthogonal planes, meaning that the model may have a higher potential to achieve better accuracy. Our model was developed to overcome the intrinsic memory requirement problem of full 3D models by integrating parameter reduction techniques such as inception blocks, dilated convolution, attention gates, and weight splitting (N-way weights). The N-way weight strategy increases the time needed for training and prediction, but enables individual model refinement through transfer learning, and allows integration of heterogeneous models. In contrast, 2.5D models require training and prediction of three models for each image plane. They can utilize a mini-batch strategy with a batch size larger than that of a full 3D model, thereby improving runtime. However, 2D or 2.5D models may lose 3D contexts, which influences parcellation accuracy.

With a classical model predicting multiple ROIs (e.g., through cross-entropy), re-weighting is rarely possible due to increased competition while minimizing the loss of multiple ROIs, where non-linear interventions occur in the model parameter space. Thus, in such cases, one should start the training from scratch, although convergence with the changed ROI configuration is not guaranteed. In contrast, competition during training never occurs with the N-way-weight strategy because weights are independent of each other.

The integration of models with heterogeneous structures is generally infeasible. We found that 3D UNet attains lower average DICEs for vessel parcellations, owing to geometrical uncertainty and randomness in shapes. With other network structures, such as partial 3D models, one may overcome uncertainty, but integration incurs re-designing and re-training of a classical model. In contrast, the N-way-weight strategy allows for individual model replacement; thus, it is possible to improve vessel parcellations without disturbing the probability map of other ROIs.

Most deep learning models for brain parcellation have been oriented towards Caucasian people [*18*]. As there are several differences in anatomy between brains of East Asian and Caucasian individuals, including shape and volume, our model can yield a better prediction, especially in the brains of older East Asian individuals, for the following reasons:

First, DeepParcellation showed robustly higher similarities in the East Asian OA group than in the other groups. That the highest DICE and lowest aHD were observed in the East Asian OA group indicates that our model is optimized for older East Asians. In the DICE evaluation, age-related dominant structural changes were observed in East Asians, although a similar number of regions showed higher DICEs between the OA and YA groups (Fig. A.1). The OA group showed higher DICEs in age-related brain regions such as the ventricles [*5*]; and WM hypointensities [*6*] compared to the YA group. Structural changes in these regions are closely associated with aging and neurodegenerative diseases, such as AD [*28*].

Second, DeepParcellation showed higher intra-subject reliability than FreeSurfer. The higher ICC of our model (0.95 vs. 0.91) indicates that DeepParcellation consistently parcellates East Asian brains. Since our model learned a global distribution of East Asian brain patterns such as shape, intensity, contrast, and volume from several thousands of East Asian samples, the predictions of unseen data are superior to those of FreeSurfer. The non-significant difference in ICCs between DeepParcellation and FreeSurfer in brains of Caucasians supports the good generalizability of the model.

Third, DeepParcellation showed better sensitivity to group differences in East Asians than did FreeSurfer. The higher statistical metrics displayed by our model were derived from significantly different mean values and variations of the normalized volume between cognitive groups (CN and AD). DeepParcellation was superior to FreeSurfer in terms of absolute Cohen’s d in selected regions (the bilateral entorhinal cortex, bilateral amygdala, bilateral hippocampus, and bilateral inferior lateral ventricles) (Fig. 3C). These regions showed significant differences in mean values and variations in comparison to FreeSurfer (Fig. 4). The difference was more robustly observed in the CN group than in the AD group because our model training was performed on more East Asian CN samples. Of particular significance is the fact that the selected regions belong to the medial temporal lobe, which is highly associated with AD [*29*]. Volume changes in these regions have been used to develop biomarkers for AD diagnosis and cognitive decline [*2*].

Finally, DeepParcellation shows a high success rate for parcellation. Some MRI data can display an abnormal intensity distribution or shape beyond the normal range of their population. Using the default command (*recon-all*), FreeSurfer can fail to process such a brain image because of inhomogeneous intensity ranges or mismatched coordinates with respect to a standard template. Contrarily, our model learned geographic patterns and image properties of brain structures from thousands of samples, rather than performing a sequence of manual algorithms. DeepParcellation, therefore, increases the chances of making valid predictions for atypical brains where FreeSurfer could fail.

Our model has some limitations that should be overcome in further studies. First, we did not train the model with different MRI acquisition parameters from multiple vendors. Most data came from our in-house GARD dataset based on the same scanner and acquisition parameters; thus, our model may have lacked technical generalization. Although our model already showed good performance on brains of Caucasians from the ADNI and OASIS databases, we think that it would be better to create a specialized model for the brains of Caucasian individuals rather than pursuing generalization, considering the anatomical differences between brains of East Asians and Caucasians. Second, segmentation failures at ROI boundaries can more severely influence smaller ROIs, which is a class imbalance problem. We may reweigh the probability values by refining some problematic ROIs before passing them to the final softmax function. Alternatively, we could replace certain predictions with newer ones by introducing a heterogeneous network with higher accuracy than that of our base 3D Attention UNet model.

Despite these limitations, DeepParcellation has great potential for use in neuroimaging studies. The predicted parcellation derived from our method can be extended to neurodevelopmental and clinical studies, such as brain age prediction or biomarker development for neurodegenerative diseases. Parcellated subcortical regions, including the putamen, caudate, and hippocampus, predict brain age with good accuracy [*30*]. Decreased cortical thickness in temporal regions was found in patients with PD [*24*] and reduced volumes in cortical regions were reported in patients with AD [*2*]. Since aging affects certain brain regions differently [*31*], accurate and precise structural measurements are critical for monitoring neurodegenerative processes. Our robust and reliable parcellation method of the brain of older individuals can guarantee higher prediction accuracy and help disease diagnosis.

The fast and robust parcellation achieved by our proposed model can accelerate big brain MRI data analysis. Our method provides crucial data for secondary applications, such as early detection or monitoring the progress of neurodegenerative diseases.

## Abbreviations

AD: Alzheimer’s disease
ADNI: Alzheimer’s Disease Neuroimaging Initiative
aHD: averaged Hausdorff Distance
CN: cognitively normal
CNN: convolutional neural network
CPU: central processing unit
DICE: dice coefficient
DKT: Desikan-Killiany-Tourville
ECB: error corrective boosting
FOV: field of view
GARD: Gwangju Alzheimer’s & Related Dementia
GM: gray matter
GPU: graphics processing unit
HD: Hausdorff distance
ICC: intra-class correlation coefficient
MRI: magnetic resonance imaging
OA: old age
OASIS: Open Access Series of Imaging Studies
PD: Parkinson’s disease
ROI: region of interest
SALD: Southwest University Adult Lifespan Dataset
TE: time of echo
TFU: Tohoku Fukushi University
TI: time of inversion
TR: time of repetition
WM: white matter
YA: young age

## Author contributions

Conceptualization: ECL, USC

Data curation: ECL

Formal Analysis: ECL, USC

Funding acquisition: KHL

Investigation: ECL, USC

Methodology: ECL

Project administration: ECL, USC, JSG

Resources: KHL, KYC, JJL, BCK, YWS, SO, ADNI

Software: ECL

Supervision: JSG, KHL, YWS, SO

Visualization: ECL, USC

Writing – original draft: ECL, USC

Writing – review & editing: ECL, USC, JSG, KHL

## Data Statement

The source codes are freely available at https://github.com/abysslover/deepparcellation. Any inquiries regarding data availability must be addressed to corresponding authors.

---

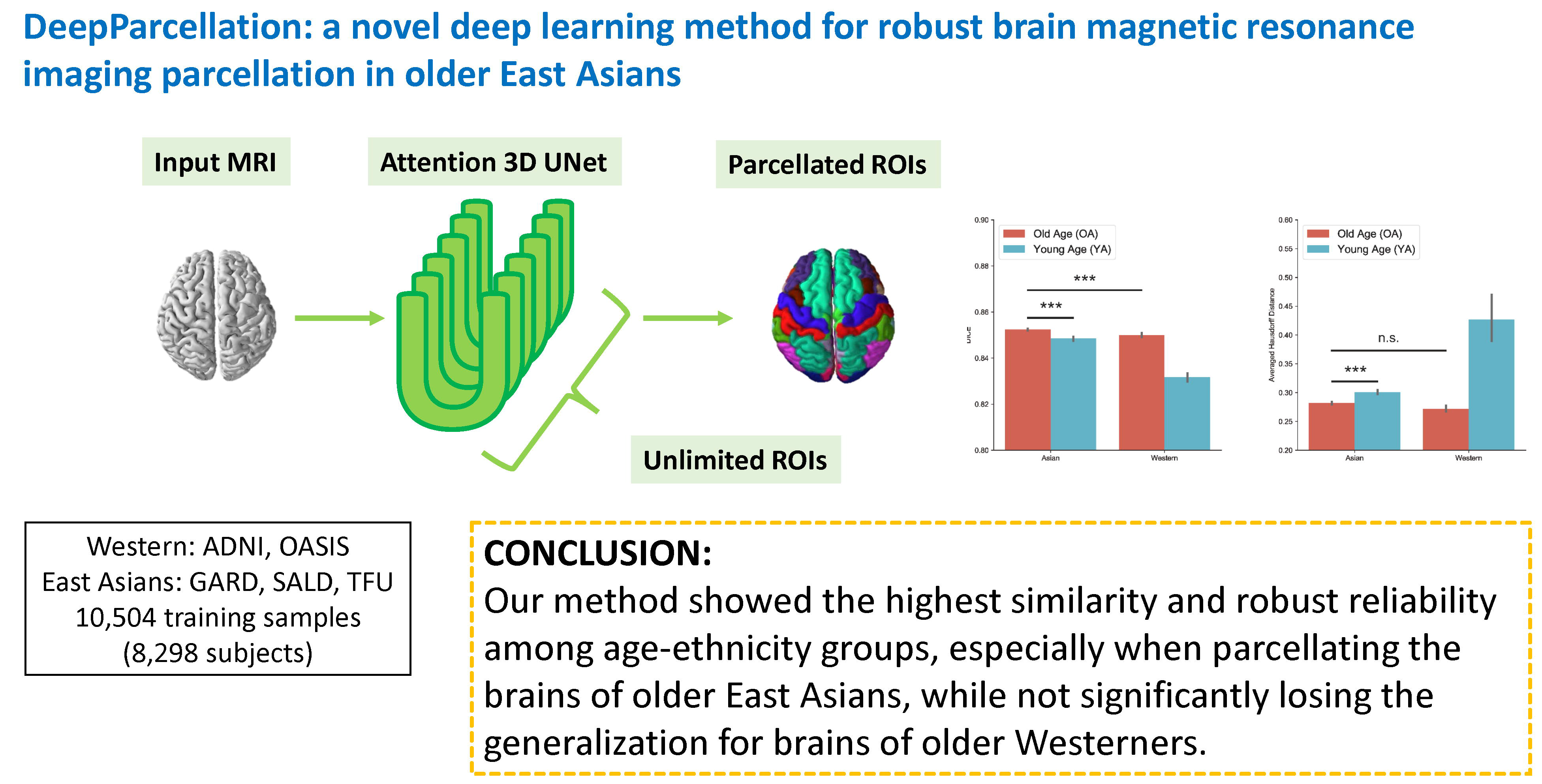

