## Supplementary Data for "DeepParcellation: a novel deep learning method for robust brain magnetic resonance imaging parcellation in older East Asians"

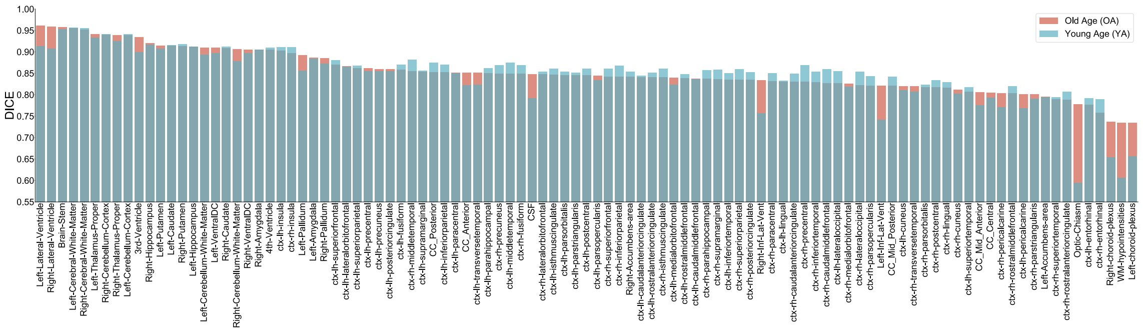


*Figure A.1 DICEs of each ROI in the age group*

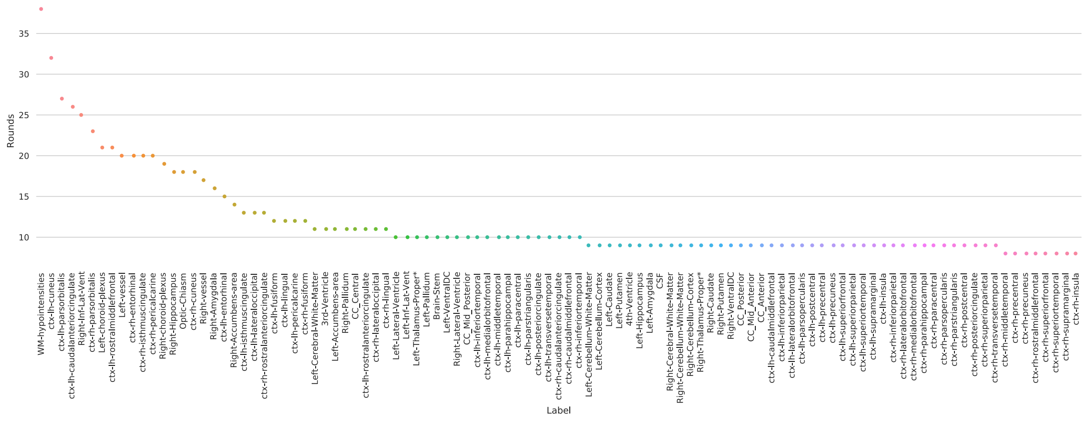


*Figure A.2 The number of training epochs. Each ROI requires the different number of epochs in order to reach at a loss convergence.*

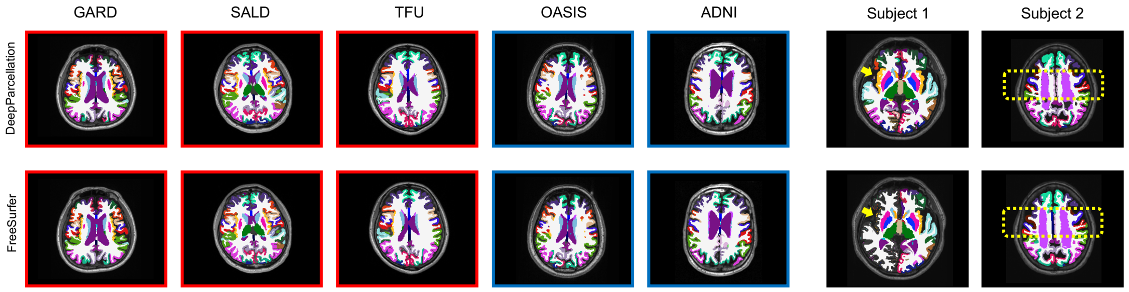


*Figure A.3 Volumetric construction of parcellated brain images from the representative subjects of different datasets (first five columns). Red and blue squares indicate Asian brains and Western brains, respectively. Examples of better parcellation of DeepParcellation compared with FreeSurfer (last two columns). (A) Failures of the right cortical parcellation in FreeSurfer (yellow arrows). (B) Wrong parcellation of right precentral and postcentral gyri in FreeSurfer (yellow dashed rectangles).*

Table A.1. Summary of deep learning models for brain MRI segmentation of a single ROI

| Name | Backend | Image Resolution | # ROIs | N (train/validation/test) | Note | Reference |
|---|---|---|---|---|---|---|
| | 3D FCNN | 160x256x256 | 1 | 15/NA/5 | Down-sampled image | [1] |
| CompNet | 2.5D custom Encoder-Decoder | 256x256 | 1 | 203/NA/203 | Tumor lesion | [2] |
| Hippodeep | 3D FCNN | 48x72x64 | 1 | 2500/NA/659 | Cropped image<br>Preprocessing required | [3] |

Table A.2. Summary of deep learning models for brain MRI segmentation of three important tissues (WM, GM, and CSF)

| Name | Backend | Image Resolution | # ROIs | N (train/validation/test) | Note | Reference |
|---|---|---|---|---|---|---|
| | 2D CNN | 256x256 | 3 | 7/NA/1 | Infant brain | [4] |
| Pyramid-LSTM | LSTM | 240x240x25 cubes | 3 | 5/NA/15 | 3 modalities<br>Extra preprocessing | [5] |
| VoxResNet | 3D CNN | 80x80x48 cubes<br>3 modalities | 4 | 25/NA/10 | Residual function<br>Multi-modality | [6] |
| MMAN | 2.5D CNN | 240x240<br>3 modalities | 3 | 4/1/15 | Dilation<br>Inception | [7] |

Table A.3. Summary of deep learning models for brain MRI segmentation of 8-30 ROIs

| Name | Backend | Image Resolution | # ROIs | N (train/validation/test) | Note | Reference |
| --- | --- | --- | --- | --- | --- | --- |
| Brainseg | 2D CNN | 256x256 | 8 | 15/NA/14 | Only works at axial axis | [8] |
| Hough-CNN | 2D,2.5D,3D CNN | 31<br>31x31<br>31x31x31 cubes | 26 | 8/NA/24 | 3D model shows better accuracy | [9] |
| Mesh-Net | 3D FCNN | 256x256x256 | 3, 8 | 20/2/100<br>5/NA/15<br>3/1/1 | Dilation<br>Unexpected artifacts | [10] |
| SD-Net | 3D UNet | 256x256x64 | 26, 24 | 581+15/5/10<br>581+10/5/5 | Deconvolution<br>FCNN<br>Error corrective boosting | [11] |
| DeepNAT | 3D CNN | 23x23x23 cubes | 25 | 20/NA/10 | Multi-task learning, hierarchical segmentation, spectral coordinates, | [12] |

| | | | | | | |
|---|---|---|---|---|---|---|
| | | | | | conditional random field | |
| 3D FCNN BrainStruct | 3D FCNN | 256x256x256 | 8 | 150/NA/923 | Extra pre-/post-processing | [13] |
| QuickNAT | 2.5D CNN | 256x256 | 27 | 581+28/NA/191 | Aggregation Median frequency balancing | [14] |
| PSACNN | 3D UNet+CNN | 96x96x96 cubes | 3,9,12 | 16/3/94 | Synthesize multi-modality images | [15] |
| SW-3D-UNet | 3D UNet | 32x32x32 cubes | 3, 25 | 4/1/15<br>20/NA/10 | Losing some 3D contexts | [16] |

Table A.4. Summary of deep learning models for brain MRI segmentation of over 50 ROIs

| Name | Backend | Image Resolution | # ROIs | N (train/validation/test) | Note | Reference |
|---|---|---|---|---|---|---|
| | 2D CNN | 28x28 + 14x14x4 patches | 56 | 20/1/19 | | [17] |
| SegNet | 2D+3D CNN | 29x29x3 patches + 13x13x13 cubes | 134 | 30/NA/5 (approx..) | | [18] |
| | 2D CNN | 25x25,51x51,75x75 patches | 134 | 10/NA/20 | Infant Losing global context | [19] |

| | | | | | | |
|---|---|---|---|---|---|---|
| HighRes3DNet | 3D CNN | 96x96x96 cubes | 155 | 443/50/50 | Dilation Residual Function | [20] |
| BrainSegNet | 2D+3D CNN | 31x31x6+21x21x21x2 cubes | 134<br>32<br>54<br>67<br>83 | 15/NA/20<br>9/NA/9<br>20/NA/20<br>10/NA/10<br>15/NA/15 | Multi-filter CNN | [21] |
| NeuroNet | 3D Encoder-Decoder | 128x128x128 | 139 | 5000/10/713 | Large-scale training | [22] |
| AssemblyNet | 3D UNet | 3x3x3x125<br>32x48x32x2 cubes | 135 | 45/NA/19 | 125 UNets<br>Extra pre/post processing | [23] |
| SLANT | 3D UNet | 86x110x78x8 cubes<br>96x128x88x27 cubes | 133 | 5111+45/NA/5+27+13 | Extra preprocessing | [24] |
| ParcelCortex | 3D UNet | 96x96x48x2 | 62<br>130<br>96 | 2300/NA/41 | Down-sampling<br>Multi-atlas hemisphere | [25] |
| FastSurfer | 2.5D | 256x256 patches | 95 | 140/20/649 | | [26] |

Table A.5. Runtime of FreeSurfer 7.1 version. We calculate the runtime of FreeSurfer using GARD cohort data including those not involved in training, and validating the DeepParcellation model. We could not clearly observe a linear relation between the runtime of FreeSurfer and the number of threads. Mean hours are denoted with a 95% confidence interval.

| # threads | # samples | Mean hours | Min. hours | Max. hours |
|---|---|---|---|---|
| 1 | 6821 | 13.22±0.07 | 5.48 | 38.35 |
| 16 | 15 | 3.56±0.18 | 3.20 | 4.55 |
| 24 | 15 | 9.4±0.14 | 9.04 | 9.89 |
| 100 | 5 | 8.43±3.49 | 6.99 | 13.44 |
| 168 | 2 | 11.14±37.29 | 8.20 | 14.07 |
| 256 | 3 | 4.98±7.09 | 3.17 | 8.27 |

Table A.6. Characteristics of training datasets.

| Dataset | Partition | # Samples | # Subjects |
|---|---|---|---|
| ADNI | Train | 808 | 764 |
| ADNI | Val | 113 | 96 |
| ADNI | Test | 110 | 96 |
| ADNI | Total | 1,031 | 956 |
| GARD | Train | 5,392 | 4,028 |
| GARD | Val | 909 | 503 |
| GARD | Test | 865 | 504 |
| GARD | Total | 7,166 | 5,035 |
| Mindboggle101 | Train | 0 | 0 |
| Mindboggle101 | Val | 0 | 0 |
| Mindboggle101 | Test | 101 | 101 |
| Mindboggle101 | Total | 101 | 101 |
| OASIS | Train | 1,263 | 1,263 |
| OASIS | Val | 158 | 158 |
| OASIS | Test | 158 | 158 |
| OASIS | Total | 1,579 | 1,579 |
| SALD | Train | 0 | 0 |
| SALD | Val | 0 | 0 |

| | | | |
|---|---|---|---|
| SALD | Test | 487 | 487 |
| SALD | Total | 487 | 487 |
| TFU | Train | 0 | 0 |
| TFU | Val | 0 | 0 |
| TFU | Test | 140 | 140 |
| TFU | Total | 140 | 140 |
| ALL | Train | 7,463 | 6,055 |
| ALL | Val | 1,180 | 757 |
| ALL | Test | 1,861 | 1,486 |
| ALL | Total | 10,504 | 8,298 |
